# The structure of phycobilisome with a bicylindrical core from the cyanobacterium *Synechococcus elongatus* PCC 7942

**DOI:** 10.1101/2025.04.28.650843

**Authors:** Zhenggao Zheng, Chengying Ma, Hongrui Wang, Guopeng Wang, Chunxia Dong, Ning Gao, Jindong Zhao

## Abstract

Phycobilisomes (PBSs) are the major light-harvesting complexes in the cyanobacteria and red algae and they consist of a central core and peripheral rods that are attached to the core. The PBS cores contain 2-5 allophycocyanin cylinders that are organized by ApcE. At the present, structures of PBS with tricylindrical and pentacylindrical cores have been determined while the structure of the PBS with a bicylindrical core is yet to be revealed. Here we report the cryo-EM structure of PBS with bicylindrical core from *Synechococcus elongatus* PCC 7942 (*Synechococcus* 7942) at 2.98-Å resolution. Similar to the PBS with a tricylindrical core, six peripheral rods are attached to the core by the rod-core linker protein CpcG in the PBS of *Synechococcus* 7942 even though the core lacks the top AP cylinder, which is important for the attachment of peripheral rods to the tricylindrical cores. We found that the C-terminus of ApcE in the *Synechococcus* 7942 was involved in interacting with both CpcG and CpcB of a top peripheral rod, compensating for the absence of the top AP cylinder of the core and maintaining PBS stability. Analysis of the bilin distribution reveals that distance of excitation energy transfer from top peripheral rods to the terminal emitters is approximately 15% shorter compared to the PBS with tricylindrical cores. Although there are 30% fewer bilin chromophores in the *Synechococcus* 7942 PBS core compared with the tricylindrical core, the aromatic residue ring in the *Synechococcus* 7942 PBS core is conserved, supporting the suggestion that these aromatic residues from AP and linker proteins are critical to the energy transfer of PBS.

## Introduction

Phycobilisomes (PBSs) are water-soluble supramolecular pigment-protein complexes located on the stromal side of the thylakoid membranes and serve as the light-harvesting antennae for photosynthesis in cyanobacteria and red algae (Adir et al. 2020; Bryant and Gisriel 2024; Sidler 1994). PBSs are composed of phycobiliproteins (PBP), which contain covalently attached phycobilins as chromophores, and linker proteins (De marsac and Cohen-Bazire 1977). The light energy absorbed by PBS is transferred to both photosystem II (PSII) and photosystem I (PSI) to drive photosynthetic electron transfer (Dong et al. 2009; Fork and Satoh 1983; Sidler 1994,). Structurally, PBSs contain two substructures: a central core, which are organized by ApcE, and peripheral rods, which are attached to the cores through linker proteins (Bryant et al. 1979; Watanabe and Ikeuchi 2013). The PBP of the central cores PBSs are allophycocyanin (AP) while the PBP in the peripheral rods are phycocyanin (PC), phycoerythrocyanin (PEC) or phycoerythrin (PE) (Sidler 1994). While five different morphological types of PBSs are found in the cyanobacteria and red algae (Bryant and Gisriel 2024), most of the cyanobacteria have the hemidiscoidal PBSs. Diversification of the core morphology in hemidiscoidal PBS are observed as their central cores could contain two (bicylindrical), three (tricylindrical) or five (pentacylindrical) AP cylinders depending on how many REP domains (Pfam00427) that the core-membrane linker protein L_CM_ (ApcE) has (Bryant and Canniffe 2018; Bryant and Gisriel 2024; Domínguez-Martín et al. 2022; Ducret et al. 1998; Glauser et al. 1992; Glazer et al. 1979; Sidler 1994; Zheng et al. 2021). PBSs with either bicylindrical or tricylindrical cores have six peripheral rods and a single CpcG is utilized for the attachment of the rods to the core while PBSs with a pentacylindrical core have eight peripheral rods and more than one CpcG linkers are utilized for their attachment to the cores (Glazer et al. 1979; Zheng et al. 2021). The cryo-EM structures of the hemidiscoidal PBSs with tricylindrical cores from *Synechococcus* sp. PCC 7002 (Zheng et al. 2021) and *Synechocystis* sp. PCC 6803 (Domínguez-Martín et al. 2022) and pentacylindrical cores from *Nostoc* sp. PCC 7120 (Zheng et al. 2021) and *Thermosynechococcus vulcanus* (Kawakami et al. 2022) have been determined. However, the structure of the hemidiscoidal PBS with a bicylindrical core has not been reported.

The majority of PBS cores contains two terminal emitters, the α_ApcE_ domain, which is a variant of ApcA and located at the N-terminus of ApcE (Adir et al. 2020), and ApcD, which is also a special ApcA variant (Ley et al. 1977). They are responsible for energy transfer from PBSs to the photosystem II (PSII) and photosystem I (PSI) (Dong et al. 2009; Zheng et al. 2025). Historically, the position of the α_ApcE_ domain and ApcD was studied by Glazer and his coworkers and it was suggested that the two terminal emitters could be located in different hexamers of the AP cylinders based on their biochemical analyses of the PBS with a bicylindrical core from *Synechococcus* sp. PCC 6301 (Bryant 1988; Lundell and Glazer 1983; Yamanaka et al. 1982). This debate was thought to be solved with the determination of the PBS structure with the cryo-EM technology (Zhang et al. 2017). The positions of the α_ApcE_ domain and ApcD in the red algal PBS are located in trimer (layer) 3 and trimer 4 of the two basal AP cylinders, respectively, i.e. the two terminal emitters are located in the same AP hexamer (Zhang et al. 2017). This arrangement of the terminal emitters is believed to be conserved in all PBSs with a cryo-EM structure (Domínguez-Martín et al. 2022; Jiang et al. 2023; Ma et al. 2020; Zheng et al. 2021). However, Kawakami et al. (Kawakami et al. 2022) reported that the cryo-EM structure of the PBS from *T. vulcanus*, in which the ApcD subunit was reportedly located in the trimer 1. Because the α_ApcE_ domain and ApcD are critical to energy transfer from PBS to the reaction centers, it is desired to determine the positions of α_ApcE_ domain and ApcD in a bicylindrical core. There are two other intriguing questions that are related to the bicylindrical core. Firstly, as the top AP cylinder serves as the attachment site for the two top peripheral rods in the tricylindrical cores (Domínguez-Martín et al. 2022; Zheng et al. 2021;), how the peripheral rods are attached to a bicylindrical core that lacks the top AP cylinder? Secondly, would the absence of the top AP cylinder of the core have any impact on the formation of the aromatic ring, which is present in all PBSs with tricylindrical and pentacylindrical cores (Ma et al. 2020; Zheng et al. 2021). Here, we report the cryo-EM structure of the PBS from *Synechococcus* sp. PCC 7942 (*Synechococcus* 7942). Our results show that the α_ApcE_ domain and ApcD are located in the same hexamer of the central core and that the ApcE is directly involved in the attachment of the peripheral rods to the bicylindrical core of the PBS.

## Results

### Structural Organization of the PBS with bicylindrical core

PBS from *Synechococcus* 7942 was found less stable compared with the PBSs with tricylindrical cores and they were isolated with sucrose density gradient centrifugation in 0.75 M K/Na-PO4 buffer containing 0.0125% glutaraldehyde to minimize PBS complex disintegration. The absorption spectrum of the isolated PBS had a smaller shoulder in 650 nm region because there is less AP in the central core as compared with the PBSs with tricylindrical and pentacylindrical cores (Fig. S1). The structure of the PBS from *Synechococcus* 7942 was reconstructed at a resolution of 2.98 Å by cryo-EM single-particle method (Fig.S2) in the presence of the cross-linker during the sample preparation. As shown in Fig. 1, the dimensions of the reconstructed PBS from *Synechococcus* 7942 are ∼480 Å in width, ∼260 Å in height, and ∼200 Å in thickness (Fig. 1a and 1b). The modeled PBS has a bicylindrical core and contains 24 αβ monomers of AP in the core, 72αβ monomers of phycocyanin (PC) in the rods, 6 rod linker proteins (L_R_), 6 rod-core linker proteins (L_RC_), and 4 core linker proteins (ApcC). Additionally, two ApcG (Residues 72-108) subunits were also partially resolved and found to bind to two central layers (layers 2 and 3) of the core (Fig.1c and 1d). The organization of the linker skeleton in *Synechococcus* 7942 PBS is similar to that found in PBSs with a tricylindrical cores except that it has only four Rep-domains (Pfam00427) in the core region (Fig. 1c and 1d).

**Fig. 1.**
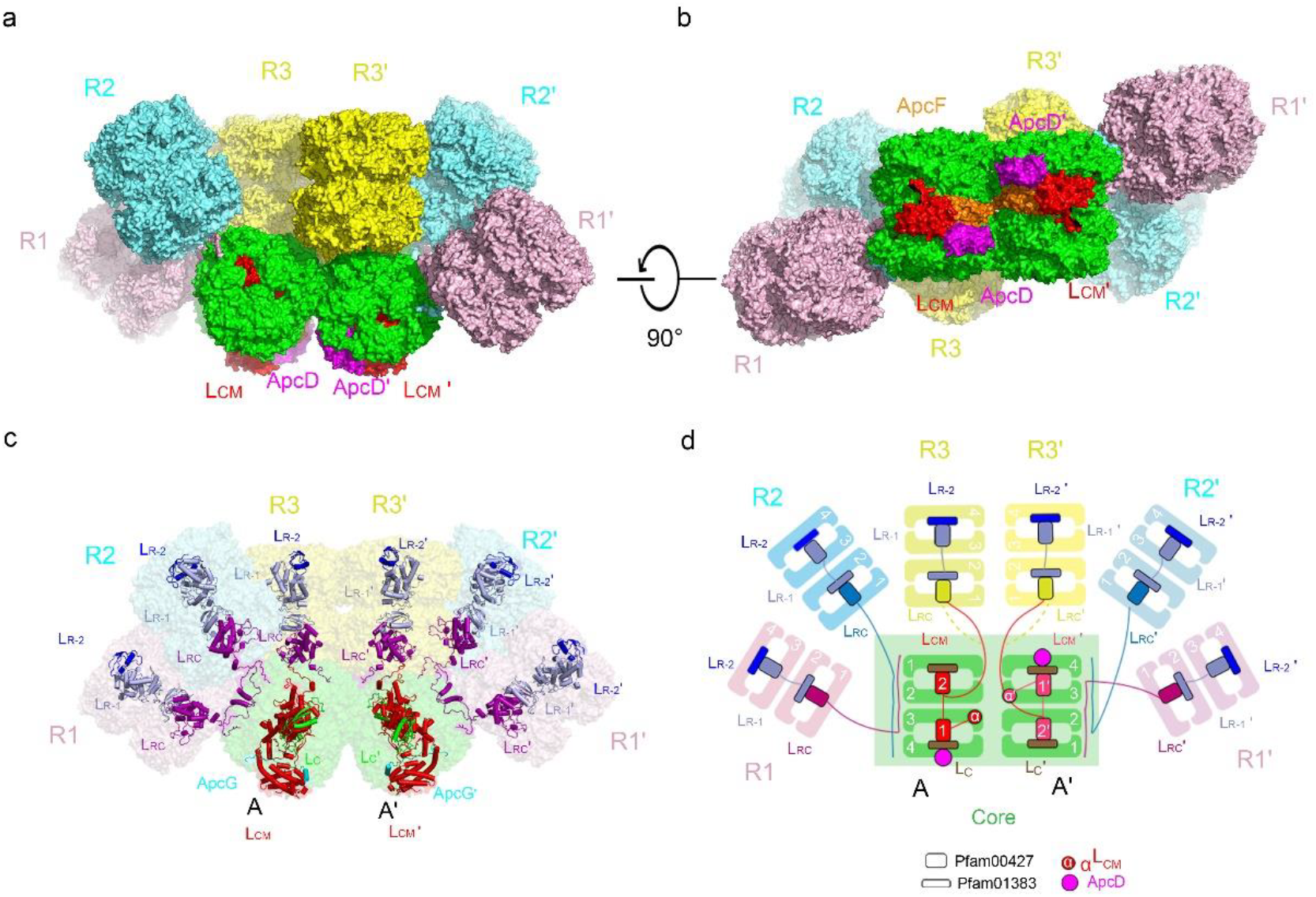
Overall structure of the PBS from *Synechococcus* 7942. **a** Side view of the PBS from *Synechococcus* 7942. The cryo-EM map is displayed in surface representation, with the core and peripheral rods colored separately. AP trimer, green; Rod1, pink; Rod2, cyan; Rod3, yellow. ApcF, ApcD, and L_CM_ are highlighted in orange, magenta, and red, respectively. **b** Bottom view of the *Synechococcus* 7942 PBS. **c** Distribution of linker proteins in the PBS from *Synechococcus* 7942. Models of linker proteins are shown in cartoon representation and colored separately. **d** Schematic models of the *Synechococcus* 942 PBS architecture. The protein linkers between core and rod are indicated with line. The linker proteins are schematically represented with rectangular boxes as indicated. The 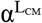 and ApcD are represented as circles and shown as indicated.

The PBS from *Synechococcus* 7942 has six peripheral rods (rods R1/R1’, R2/R2’, and R3/R3’) and each rod contains two hexamers in the model (Fig. 1). The bicylindrical core consists of two AP cylinders (A/A’) arranged antiparallelly. The α_ApcE_ domain of ApcE is located in trimer (layer) 3 while ApcD is located in trimer 4 of the two AP cylinders. The spatial positions of ApcE, ApcD and ApcF are identical in PBSs with bicylindrical, tricylindrical and pentacylindrical core. Therefore, although the AP core exhibits significant structural diversity, the component and basic structural framework of the cores are evolutionarily conserved.

### Linker proteins

The PBS assembly relies on four types of linker proteins: rod linker proteins (CpcD, L_R_), rod-core linker protein (CpcG, L_RC_), core linker protein (ApcC), and the core-membrane linker protein (ApcE, L_CM_). In *Synechococcus* 7942 PBS, six L_RC_ copies adopt three distinct C-terminal extensions (CTE) conformations to accommodate structural variations of the core (Fig. 2a-d). CTE from L_RC_ of rod-1 (R1) traverses core layer 3 to layer 1, while CTE from L_RC_ of rod-2 (R2) traverses core layer 2 to layer 4. The CTEs of the L_RC_ in R3/R3′ lacked sufficient density for modeling (Fig. 2b) likely due to the high flexibility of the top peripheral rods. Structural alignment of L_RC_ linkers from *Synechococcus* 7002 (tricylindrical PBS) and *Synechococcus* 7942 highlighted CTE conformational plasticity, contrasting with conserved Pfam00427 domains (Fig. 2e). The rod linker L_R_ exhibited high conservation between species as can be observed with aligned Pfam001383 and Pfam00427 domains (Fig. 2f).

**Fig. 2.**
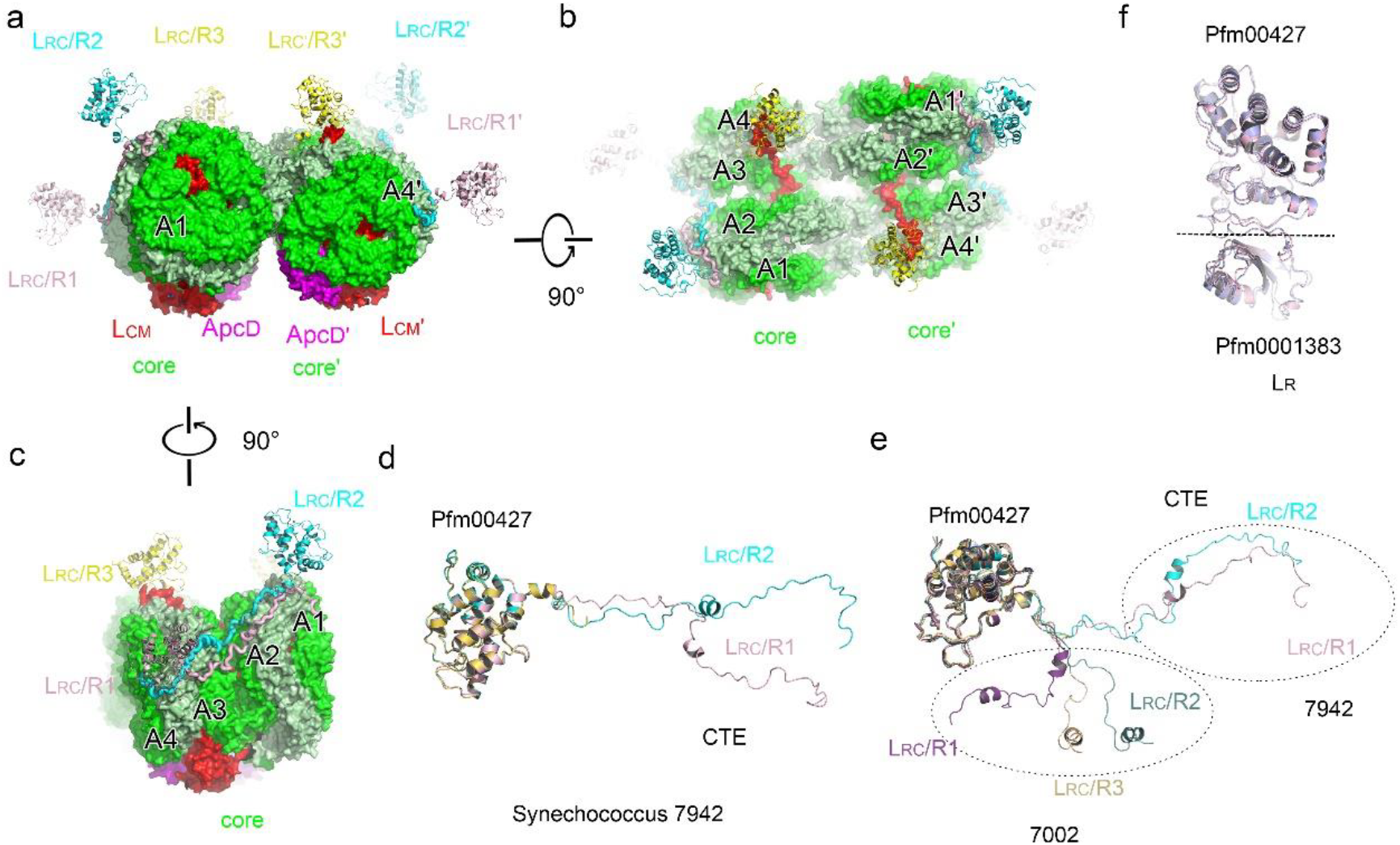
Structural comparison of linker proteins. **a** Spatial distribution of L_RC_ proteins relative to the PBS core of *Synechococcus* 7942. The L_RC_ proteins and the core cylinders are shown in cartoon and surface representation, respectively. **b**,**c** A 90°-rotated view compared to (**a**) **d** Structural comparison of L_RC_ proteins from *Synechococcus* 7942. **e** Spatial distribution comparison of L_RC_ proteins from *Synechococcus* 7942 and *Synechococcus* 7002 (PDB:7EXT). The L_RC_ proteins from 7942 and 7002 are shown as indicated. **f** Structural comparison of L_R_ protein from *Synechococcus* 7942 and *Synechococcus* 7002. The L_R_ proteins from 7942 and 7002 are shown in pink and light purple, respectively.

The PBS core is organized by the core-membrane linker L_CM_, or ApcE, which has one α^ApcE^ domain followed by two repetitive (REP) Pfam00427 domains (Fig. 3a). Comparison of these domains of the ApcE from *Synechococcus* 7942 and *Synechococcus* 7002 by superimposition (Fig. 3b-d) shows that these domains are very similar conformations and spatial arrangement. However, the third REP domain present in the *Synechococcus* 7002 ApcE (Fig. 3b), which is responsible for organizing the top AP cylinder of the core, is absent in the ApcE of *Synechococcus* 7942. As a result, the *Synechococcus* 7942 PBS core lacks the top AP cylinder. Since the top AP cylinder of the core is crucial for the attachment of the two top peripheral rods in the tricylindrical cores, the mechanism for the attachment of the two top peripheral rods to the bicylindrical core would be different. Indeed, the cryo-EM structure of the *Synechococcus* 7942 PBS reveals that the resolved C-terminus of ApcE in the *Synechococcus* 7942 PBS interacts with both L_RC_ and CpcB of a top peripheral rod. The C-terminus of ApcE corresponds to the loop region between the second Rep-domain and the third Rep-domain of ApcE in PBS with tricylindrical or pentacylindrical core. The Q696 and R699 from ApcE protein interact with N112 and E116 from CpcB, respectively. The E701 and R704 from ApcE interact with T172 and K177 from CpcG, respectively. The R705 from ApcE interacts with D115 from another CpcB (Fig. 3e). These interactions play important roles in rod-core association and therefore compensate for the absence of the top AP cylinder of the core and allow the attachment of peripheral rods to the core from above (Fig.3e).

**Fig. 3.**
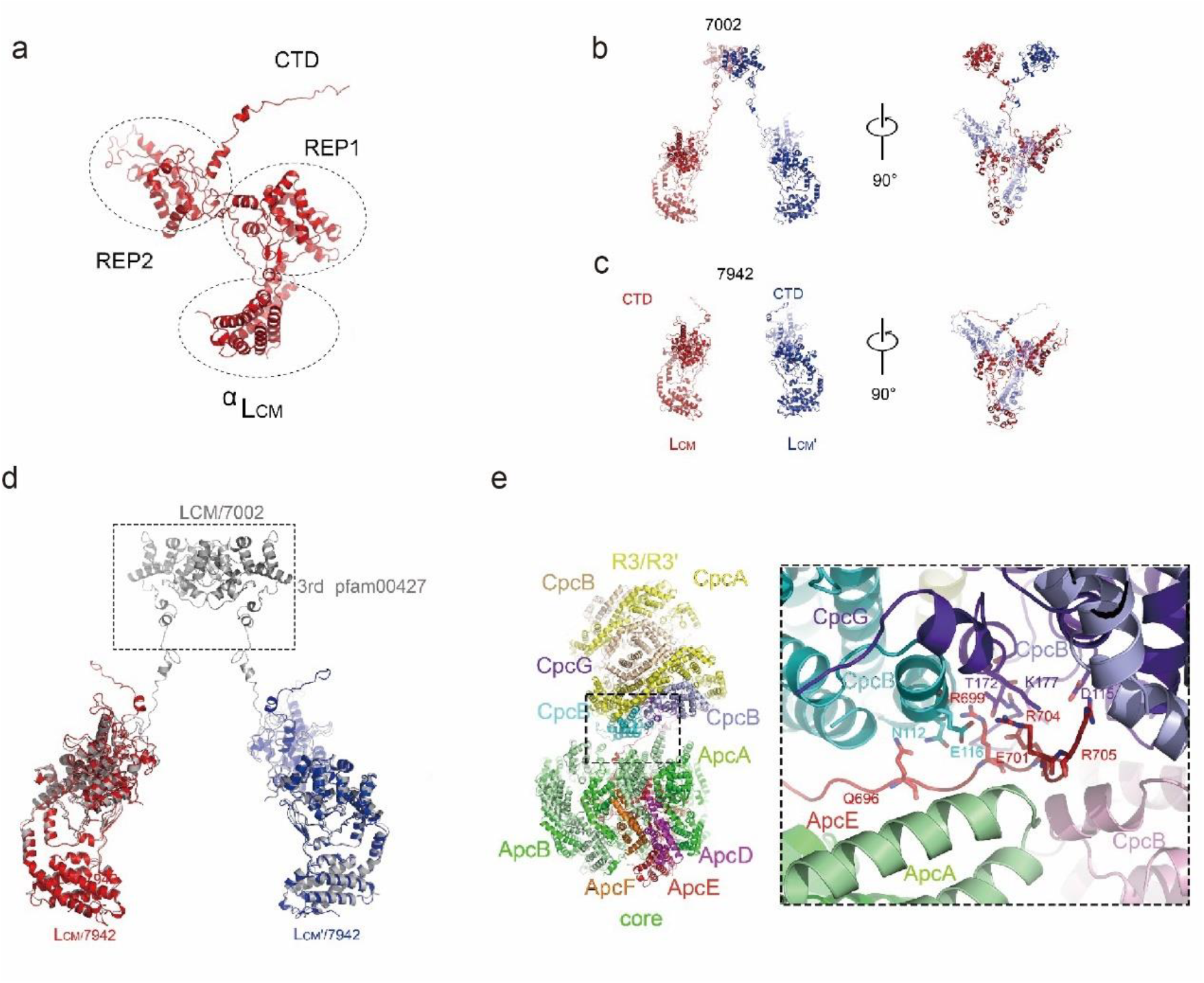
Structural comparison of ApcE. **a** The domain organization of ApcE. Each domain of ApcE is shown as indicated. **b-d**, Structural comparison of ApcE from *Synechococcus* 7942 and *Synechococcus* 7002 (PDB:7EXT). The ApcE from *Synechococcus* 7002 are shown in grey. The ApcE of two cores from *Synechococcus* 7942 are shown in red and blue, respectively. **e** The overall structure view of core A and Rod3. The enlarged view shown of detail interaction of CTE from ApcE and Rod3 (right). The Q696 and R699 from ApcE protein interact with N112 and E116 from CpcB, respectively. The ApcE, ApcD, ApcA, and ApcB from the core are shown in red, magenta, green, and pale green, respectively. The CpcG, and CpcA, CpcB from layer 2-4 are shown in purple, yellow and wheat, respectively. The CpcB from layer 1 are shown in pink, cyan, and light purple, respectively.

### Bilins distribution in PBS core

Figure 4a illustrates the spatial distribution of bilins within the PBS of *Synechococcus* 7942. We identified a total of 264 phycocyanobilin (PCB) in the model of the *Synechococcus* 7942 PBS with 48 PCB in the core and 216 PCB in the rods (Fig. 4a). The average distance for excitation energy transfer (EET) from top two peripheral rods to the terminal emitters in PBS with a bicylindrical core is smaller than that in PBS with tricylindrical cores. The most probable paths of EET to the terminal emitters (ApcD and the α^ApcE^ domain) are deduced based on the shortest distances among the bilin pairs (Fig. 4b-g). In *Synechococcus* 7942 PBS, energy from R1 could be transferred to ApcD and α^ApcE^ domain through trimer 4 and trimer 3 in the AP cylinder, respectively. Energy from R2 is more likely transferred to α^ApcE^ domain while energy from R3 might be transferred to ApcD and α^ApcE^ domain through two separate pathways (Fig. 4b-d). In all the predicted pathways mentioned above, a key step is EET from peripheral rods to the core. Like all other hemidiscoidal PBSs, the distances from the core-proximal bilins in rods to the nearest bilins in the core are in a range from 37 Å to 42 Å in *Synechococcus* 7942 PBS (Fig. 4e-g).

**Fig. 4.**
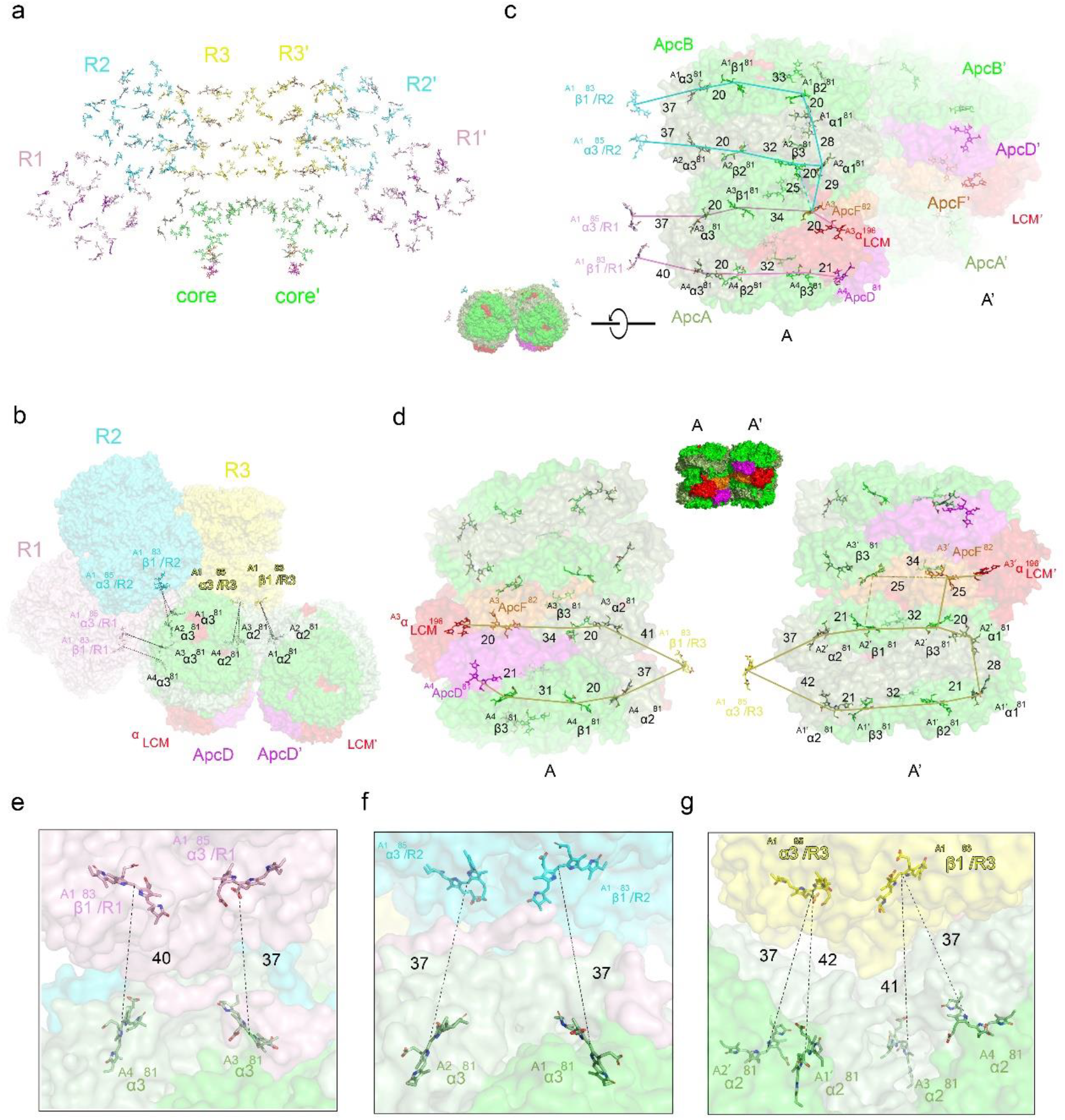
Key bilins and possible pathways for energy transfer in the core. **a** The bilin distribution in the PBS from *Synechococcus* 7942. All bilins are shown in stick representation and color-coded according to their spatial locations. **b-d** Different view shown of the core with possible energy-transfer pathways in the PBS of *Synechococcus* 7942. ApcE, ApcA, ApcB, ApcD, and ApcF are colored red, pale green, green, magenta, and orange, respectively. Bilins in rods and core are painted the same colors as the corresponding proteins. **e-g** The detail plausible energy transfer paths from bilins on the rod to the core, with distances indicated.

The interaction between bilins and the surrounding amino acid residues from PBP and the linker proteins is critical to EET (Zhang et al. 2017; Zheng et al. 2021). We analyzed the spatial distribution of the residues that could interact with bilins and influence EET in PBS cores. The formation of AP trimer establishes a conserved circumferential arrangement of aromatic residues (Fig. 5a,b). The aromatic residues of the two REP domains of ApcE, which are located within cavities of AP hexamers, also interact with the bilins in the core. Notably, the Rep domain in REP1 of ApcE located in the A3/A4 hexamer cavity contributes a rich spatial distribution of aromatic residues (Fig. 5c). These aromatic residues are cooperatively assembled into an extended π-conjugated network. The Pfam00427 domain from CpcG contains conserved aromatic residues, and the distribution of these aromatic residues are polarized around the bilins (bilins β1 and α3, Fig. 4) of the core-proximal hexamers of the peripheral rods (Fig. 5d). Although the *Synechococcus* 7942 PBS core contains 30% fewer bilin chromophores than a tricylindrical core, the aromatic residue ring in the *Synechococcus* 7942 PBS core is conserved, supporting the suggestion that these aromatic residues from AP and linker proteins are critical for energy transfer in PBS.

**Fig. 5.**
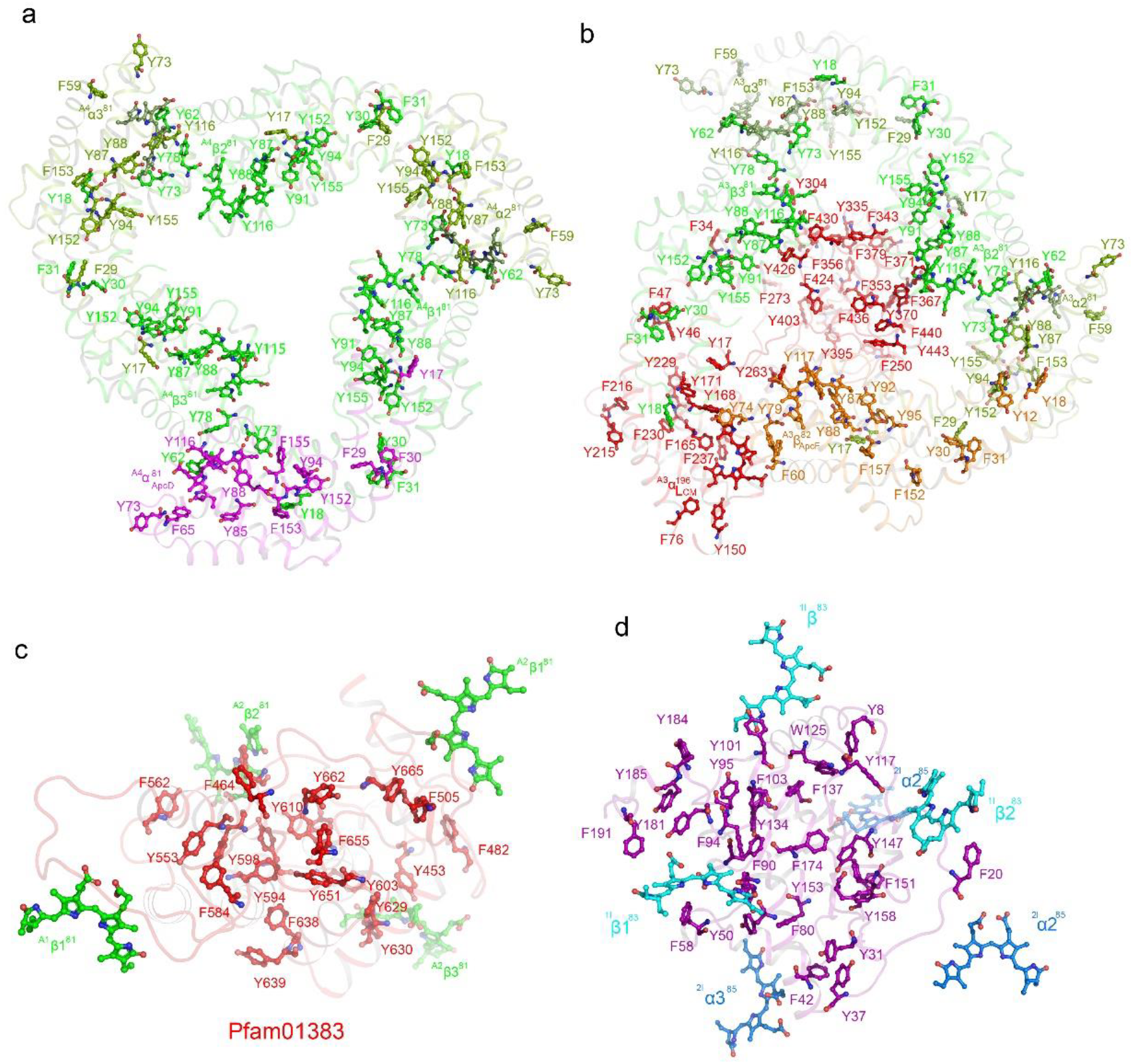
The spatial distribution of aromatic residues. **a** The spatial distribution of aromatic residues in layer A4 of cylinder. **b** The spatial distribution of aromatic residues in layer A3 of cylinder. **c** The spatial distribution of aromatic residues of the Pfam00427 domain in the REP2 region of ApcE. The ApcE, ApcD, ApcA, and ApcB are shown in red, magenta, green, and forest green, respectively. The bilins from ApcB are colored green. **d** The spatial distribution of aromatic residues of the Pfam00427 domains of CpcG. The CpcG is shown in purple. The bilins from ApcB are shown in cyan and blue, respectively.

## Discussion

Much of our knowledge about PBS came from studies of the PBS with bicylindrical core from *Synechococcus* 6301 by Glazer and co-workers (Glazer 1985 and Glazer 1989). On the other hand, the structure of the PBS with a bicylindrical core has not been reported when the cryo-EM structures of nearly all other types of PBSs were reported (Bryant and Gisriel 2024). An important issue that was not resolved by the early studies is the location of ApcD relative to the location of α^ApcE^ domain (Bryant 1988). The results from biochemical analysis could not determine if ApcD was in the same AP hexamer with α^ApcE^ domain or they are located in separate AP hexamers (Bryant 1988). This issue was thought to be resolved when cryo-EM structures of the PBSs from red algae (Ma et al. 2020; Zhang et al. 2017) and cyanobacteria (Domní guez-Martní et al. 2022; Zheng et al. 2021) were obtained and they showed that ApcD and α^ApcE^ domain were located in the layer 4 and 3, respectively, i.e. they are located in the same AP hexamer. However, Kawakami et al. reported that ApcD and α^ApcE^ domain were located in different AP hexamers in *T. vulcanus* PBS (Kawakami et al. 2022), raising a question if the location of ApcD in PBS cores is conserved. In this study, we determined the cryo-EM structure of the PBS from *Synechococcus* 7942, which shows that the location of ApcD and α^ApcE^ domain are the same as all other PBSs except for the PBS from *T. vulcanus*. Because α^ApcE^ domain and ApcD are critical to energy transfer from PBS to PSII and PSI, respectively (Dong et al. 2009), it is predicted that their locations in the PBS cores should be conserved. The reason for the location of ApcD in layer 1 in *T. vulcanus* PBS core is unclear (Bryant and Gisrie 2024), but it could be a special adaption of the cyanobacterium grown at high temperature.

The *apcE* gene of *Synechococcus* 7942 encodes an ApcE with only two Rep-domains, leading to the formation of PBS with a bicylindrical core, i.e. the *Synechococcus* 7942 PBS contains fewer AP cylinders in the core in comparison to all other PBSs. Several unique features are observed in the PBS with a bicylindrical core. The less AP content of the core leads to a reduced absorption in red light (Fig. S1) while its relative absorption of green light increased. Actually, a similar situation was found in the block-type PBS (Zhang et al. 2017) and hemiellipisoidal PBS (Ma et al. 2020) of red algae. Although these two types of PBSs have a tricylindrical core, the lengths of the AP cylinders are reduced: the top cylinder contains only two trimers while the basal two cylinders contain three trimers each. The net AP content in these two types of PBSs is therefore identical to that of the PBS with bicylindrical core. These types of PBSs would have an adaptive advantage for the cyanobacteria and red algae grown in an environment rich in green light or blue-green light (Zhang et al. 2017). Secondly, the lack of the top cylinder in a bicylindrical core provides a challenge for attachment of the peripheral rods as it serves as the attachment sites for the rods in tricylindrical and pentacylindrical cores. Our results show that the ApcE C-terminus compensates for the loss of the top cylinder by direct interaction with CpcG/CpcB, a mechanism that is not found in other types of PBSs. This adaptation is likely critical to the stability of rod-core connection and maximizes light absorption cross-sections by maintaining six rod attachments to the bicylindrical cores.

Another consequence of bicylindrical core is that the PBS no longer has the binding site for the orange carotenoid protein (OCP) as OCP requires tricylindrical core interactions (Domínguez-Martín et al. 2022). OCP is a key player for non-photochemical quenching (NPQ) in cyanobacteria and it is important to the cyanobacteria in response to light stress (Cogdell and Gardiner 2015; Kerfeld and Sutter 2024; Sedoud et al. 2014). In fact, the gene coding OCP is not present in *Synechococcus* 6301/7942 with bicylindrical cores while the genes encoding OCP are found in nearly all cyanobacteria, including *Gloeobacter violaceus* sp. PCC 7421, the oldest cyanobacterium, suggesting that the *ocp* genes were lost during evolution because the OCP binding sites were lost in bicylindrical core.

## Materials and Methods

### Isolation of PBS

*Synechococcus* 7942 was cultured in BG11 liquid media bubbled with 1% CO_2_ at 25°C with light intensity of 50 μmol photons m^−2^ s^−1^.. PBSs was prepared according to Zheng et al with some modifications. All operations were performed at 22 °C. All operations were performed at 22°C. Briefly, the cells were broken in 0.75 M K/Na-PO4 buffer at pH 7.0 with 1 mM phenylmethylsulfonyl fluoride by passing through three times of a French press at 20,000 psi. After 0.5 h of incubation with 1.0% n-dodecyl-β-d-maltoside (β-DDM) (w/v), samples were centrifuged at 20,000×g. The supernatant was loaded immediately onto a sucrose gradient. The sucrose gradients were made from buffer A (0.75 M K/Na-PO4 buffer, 0.0125% glutaraldehyde (w/v), pH 7.0) by adding sucrose to these concentrations: 0.25, 0.4, 0.55, 0.7, 0.85, and 1.0 M. The samples were ultracentrifuged at 35 000 rpm for 6 h using a Ti-40 rotor on Beckman Optima L-80XP centrifuge and resulted in a visible band of PBSs as the main layer of intact PBSs. The sucrose of purified intact PBSs was removed by ultrafiltration with 30 KD Millipore centrifugal filters.

### Cryo-EM sample preparation and data collection

Before preparing grids for cryo-EM, PBS samples were concentrated to ∼1.0 mg ml^-1^. Aliquots (3.5 μl) of samples were loaded onto glow-discharged (30 s) holey carbon films copper grids (Quantifoil R1.2/1.3, +2 nm C membrane, Cu 300 mesh) and waited for 45 s. 3.5 μl aliquot of a buffer (50mM Tris pH 7.0) was then added twice to the grid and mixed immediately with the sample before vitrification (to dilute the high concentration of phosphate salts). After blotting for 2 s, the grid was plunged into liquid ethane with an FEI Vitrobot Mark IV (18 °C and 100% humidity). Cryo-grids were first screened in a Talos Arctica operated at 200 kV (equipped with an FEI CETA camera). Grids of good quality were transferred to an FEI Titan Krios operated at 300 kV with Gatan BioQuantum GIF/K3 direct electron detector for data collection.

Movies were recorded with a BioQuantum GIF/K3 direct electron detector (Gatan) in the super-resolution mode at a nominal magnification of 81,000 x, with an exposure rate of 17.9 e-/Å2 per second using the EPU software. A GIF Quantum energy filter (Gatan), with a slit width of 20 eV was used at the end of the detector. The defocus range was set from-1.0 to-1.8 μm. The total exposure time was 3.84 s and 32 frames per image were acquired with an total electron exposure of ∼60 e−/ Å2. Statistics for data collection are summarized in Table S1.

### Image processing

To determine the 3D structure of the PBS with bicylindrical core, two batches of movie stacks were recorded. Raw movie frames were aligned and averaged into motion-corrected summed images with a pixel size of 1.07 Å by MotionCor2. The Gctf program (v1.06) was used to estimate the contrast transfer function (CTF) parameters of each motion-corrected image. All the following data processing was performed with Relion-3.1. 1,350 particles were manually picked and subjected to 2D classification to generate templates for automatic particle picking. Then autopicking was done to the images that were manually selected for treatment. The picked particles were subjected to one round of 2D classification, and high-quality particles were further selected for subsequent 3D classifications. Initial model was generated from these particles. After multiple rounds of 3D classification, ralatively homogeneous particles were selected for 3D refinement, resulting in a map with a 2.64 Å overall resolution after mask-based post-processing, based on the gold-standard FSC 0.143 criteria. The local resolution map was analyzed using ResMap and displayed using UCSF Chimera. Workflow of data processing was illustrated in the Fig. S2.

### Model building and refinement

The atomic models of all Synechococcus 7942 proteins were adopted from AlphaFold databases (Jumper et al. 2021). All structures were firstly docked into the density map using UCSF chimera (Pettersen et al. 2004) and PYMOL and then manually adjusted in COOT (Emsley et al. 2010). The final atomic models were refined in real space using PHENIX (Afonine et al. 2018) with secondary structure and geometry restraints applied. The final atomic models were evaluated using Molprobity (Chen et al. 2010) and the statistics of data collection and model validation were included in Table S2.

## Data availability

All data is available from the corresponding author.

## Acknowledgements

This work was supported by National Natural Science Foundation of China (32070203) to J. Z., Qidong-SLS Innovation Fund (202001539) and a special fund from Peking University to J.Z. (7101303501). We thank the Core Facilities of School of Life Sciences, Peking University for assistance with negative-staining electron microscopy and the High-performance Computing Platform of Peking University for technical help in computation. We also thank the National Center for Protein Sciences at Peking University for technical support.

## Author contributions

N.G., and J.Z. conceived the project; N.G., and J.Z. designed research; Z. Z. and H.W. isolated PBS and performed biochemical analysis; Z. Z. and H.W. performed cryo-EM experiment; C.M. and G.W. processed the cryo-EM data; C.M. performed model building for structure determination; N.G., and J.Z. wrote the manuscript with help from all authors. All authors reviewed the manuscript.

## Figure Legends

**Fig. S1.**
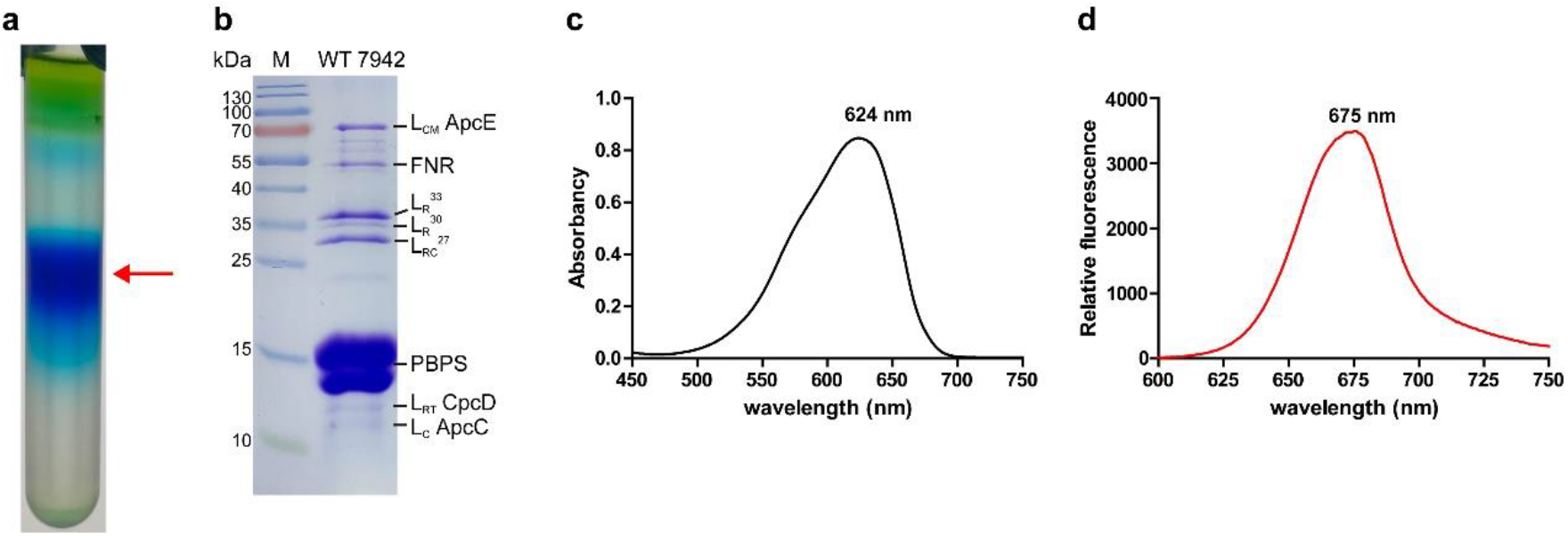
Preparation and characterization of the PBS from *Synechococcus* 7942. **a** Isolation of phycobilisomes from *Synechococcus* 7942 using sucrose density gradient centrifugation. The samples of the band with red arrow pointing were used for cryo-EM single particle analysis in this study. **b** SDS-PAGE analysis of protein components of the PBS from *Synechococcus* 7942. The gel was stained with Coomassie brilliant blue. Experiments were repeated more than three times with similar results. **c** Absorption spectrum of the PBS from *Synechococcus* 7942. **d** Fluorescence emission spectra of the PBS from *Synechococcus* 7942, excited by 590 nm at room temperature.

**Fig. S2.**
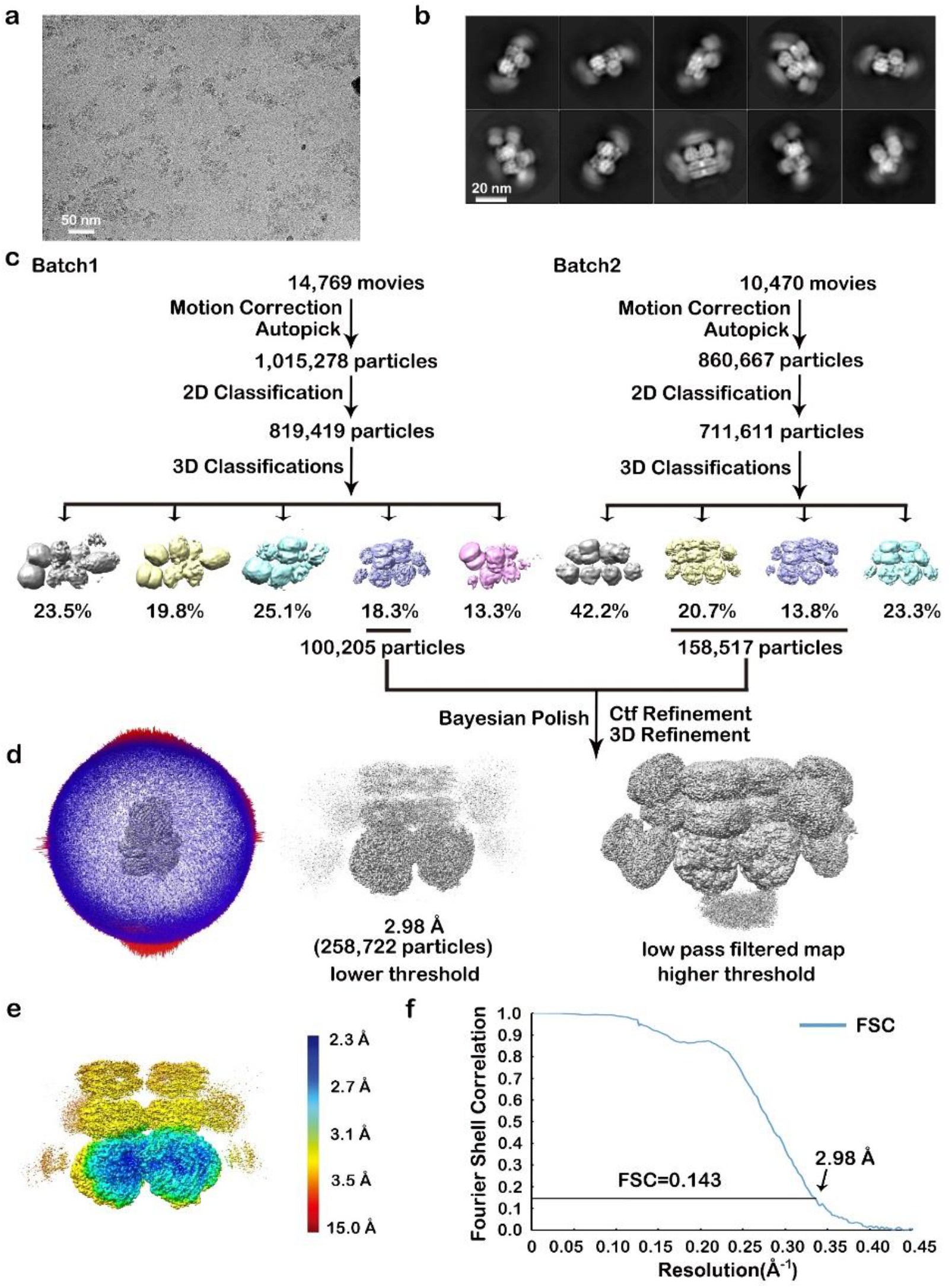
Workflow of the cryo-EM image processing. **a** A representative raw cryo-EM image. **b** Representative 2D class averages of the PBS particles from *Synechococcus* 7942. **c** Image processing workflow, including 3D classification, structural refinement, masked-based refinement, CTF refinement and Bayesian polishing. **d** Angular distribution of the PBS particles in the final round of 3D refinement. **e** Local resolution estimation of the final density map. **f** Gold-standard Fourier shell correlation (FSC) of the final cryo-EM map.

**Fig. S3.**
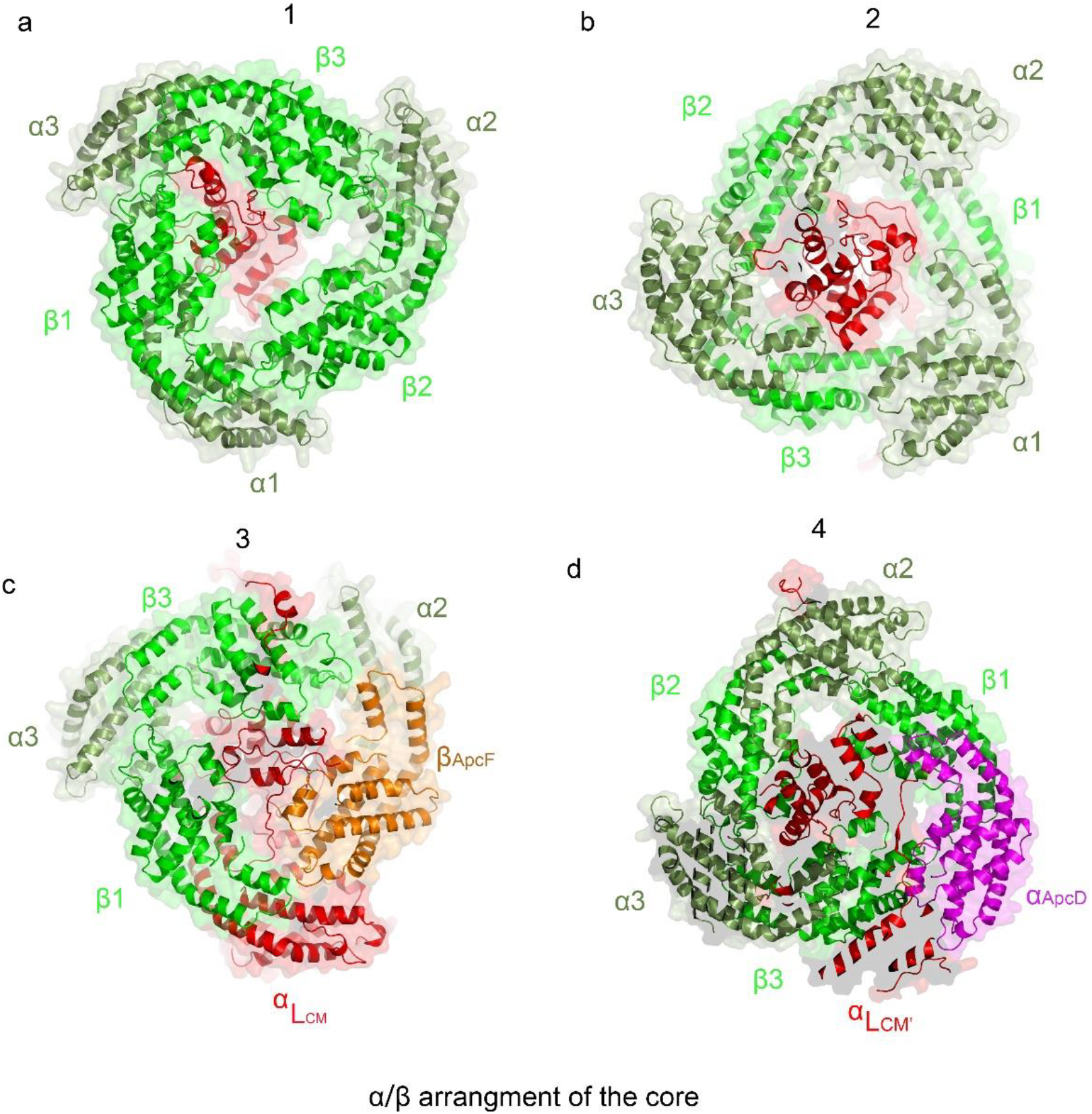
The protein arrangement of the different layers of the core. The ApcE, ApcD, ApcA, and ApcB proteins are shown in red, magenta, green, and forest green, respectively.

## Notes

### Competing Interest Statement

The authors have declared no competing interest.

